# Neural correlates of cardiac interoceptive accuracy across development: implications for social symptoms in autism spectrum disorders

**DOI:** 10.1101/630343

**Authors:** Michelle D. Failla, Lauren K. Bryant, Brynna H. Heflin, Lisa E. Mash, Kim Schauder, Samona Davis, Madison B. Gerdes, Amy Weitlauf, Baxter P. Rogers, Carissa J. Cascio

**Author notes:** **Corresponding Author:** Carissa J. Cascio, Vanderbilt University Medical Center.

## Abstract

**Background:** Interoception involves the processing of sensory information relevant to physiological functioning and is integral to building self-awareness, emotional states, and modulating social behaviors. With the role of interoception in emotional processing and social functioning, there is growing interest in characterizing interoception in autism spectrum disorder (ASD), yet, there are mixed results regarding cardiac interoceptive accuracy in ASD.

**Methods:** In this study, we explored the neural basis of cardiac interoception using an fMRI heartbeat counting task in order to assess neural correlates of primary interoception. We predicted that interoceptive-specific response in the insula, a “hub” for interoception, would be related to ASD symptomatology. We investigated the relationship of insula responses during cardiac interoceptive accuracy and a self/caregiver-reported autism-related symptom scale (Social Responsiveness Scale-2 (SRS)). Participants included 46 individuals with autism spectrum disorder (ASD) (age 8-54, mean= 19.43±10.68 years) and 54 individuals with typical development for comparison (TC, age 8-53, mean= 21.43±10.41 years).

**Results:** We found no significant difference in cardiac interoceptive accuracy or neural response to cardiac interoception in ASD. Several insula sudivisons had a curvilinear relationship to age, peaking in early adulthood. Interoceptive-specific insula response was associated with adult self-report SRS scores; this association differed by diagnostic group and self/other report.

**Conclusions:** This work suggests that 1) there is no global deficit in interoception in ASD, but that integrating interoceptive cues with social information may distinguish individuals with ASD, and 2) there is a developmental trajectory for interoceptive processing in the insula that may be relevant for socio-emotional health.

## Introduction

The human experience of emotion begins in the body, with sensory cues generated by autonomic responses to emotionally-salient events, such as a racing heart after a near-miss in traffic, or the feeling of butterflies in the stomach on a first date. This internally-generated landscape of sensations is known as interoception, though this term is now often used more broadly to describe all somatic input that confers information about the physiological state of the body, whether arising from the viscera or from the skin (1). Higher-order representations of emotional states are built from these bodily sensations; thus accuracy and interpretation of interoceptive signals can influence emotional processing and regulation (2, 3). By learning to associate interoceptive information with external stimuli, we build a sense of self in relation to others that serves as a foundation for social function(4).

While few experimental studies have investigated interoception across development (for a review see (5)), the available evidence distinctly indicates a trajectory of interoceptive perception that changes with development. Maister and collegues demonstrated that infants as young as five months can distinguish stimuli presented asynchronously vs. synchronously with their own cardiac rhythm. This implicit perception of cardiac interoceptive cues was altered by emotional state, providing support for the idea that interoception is central to the development of socio-emotional awareness (6). Older children show similar levels of cardiac interoceptive accuracy to adults in a experimental heartbeat counting paradigm (7, 8), suggesting early maturation of explicit interoceptive abilities. These studies indicate perception and processing of interoceptive cues begins early in development, thus, understanding their trajectory will be important for characterizing the role of interoception in socio-emotional development.

Given the role of interoception in emotional processing and social functioning, there is a growing interest in characterizing interoception in autism spectrum disorder (ASD) (9–11). ASD is characterized by social communication difficulties, restricted and repetitive behaviors, and altered sensory perception and reactivity. Thus, altered interoception could be related both to perceptual differences and social-emotional differences in this population. There is mounting evidence of altered insular connectivity in ASD across development (12, 13). Examining interoceptive abilities in ASD, specifically across the lifespan, may provide significant insight into the social communication deficits that are central to ASD (14, 15). Some studies report diminished cardiac interoceptive accuracy in both school-age children (16) and adults with ASD (17, 18), but most have found no significant differences (10, 11, 19–21). For a review, see (22)). While the field may be moving towards an understanding that cardiac interoceptive accuracy is not systematically/dramatically different in ASD, how these cues are interpreted, and relayed to the rest of the brain – and how they relate to autism symptoms - remains to be seen.

Understanding the role for interoceptive processing in autism related-symptoms is likely tied to one’s ability to pair interoceptive information with external events. Altered integration of interoceptive information could have early consequences. Recently, Quattrocki and Friston proposed that the ability to pair interoceptive signals, like satiety, with external cues, like a caregiver’s presence, could influence infant social learning and attachment (15). This relationship between social functioning and interoception is maintained over the lifespan. In adolesence, heightened social sensitivity increases the influence of social context on interoceptive functioning (23, 24). In adults, heightened interoceptive accuracy may be a protective factor against social exclusion (25). Given the role of social functioning in typical development, and in developmental disorders like autism spectrum disorders (ASD), it is important to understand interoceptive functioning across development and how it may influence socio-emotional health at different ages.

Examination of neural correlates of interoception provides an opportunity to assess the connection between perceptual and social-affective brain networks, while avoiding potential confounds related to self-report and experimental task performance. A meta-analysis of cardiac interoceptive fMRI studies shows consistent response in the somatomotor cortex, medial frontal regions including the cingulate, and most prominently, in the bilateral insula (26). The posterior insula contains primary interoceptive cortex (27), while salience and emotional processing involves the anterior insula (28). Studies have also shown a specific role of the anterior insula in modulating the influence of interoceptive processing on social behaviors (29, 30). While there has been limited work specifically addressing developmental changes in interoceptive neural responses, there is evidence of a non-linear developmental trajectory, with peak neural responses (24) and structural changes in insular connectivity (31) emerging in adolescence/early adulthood.

In this study, we aim to examine the neural responses underlying unisensory cardiac interoceptive processing in ASD and to understand how interoceptive processing may change across development. By investigating how interoception changes with age, we can better understand its role in both typical and atypical social development. The use of neuroimaging provides a window into stages of perceptual processing that *precede* interoceptive awareness and its influence on socio-emotional functioning. Thus, in the current study, we used functional neuroimaging in combination with a heartbeat counting task (Schandry, 1981). Among several alternatives, we chose this task for two reasons: 1) its widespread use maximizes the ability to interpret findings in the context of a large literature on this topic, and 2) its interoceptive “purity”—compared to other established interoceptive tasks like the heartbeat discrimination task that requires comparison of external stimuli to cardiac rhythms, the Schandry task demands focus on interoceptive signals alone. This feature allows for the establishment of neural correlates of interoception with minimal exteroceptive influences.

Given the mixed results regarding cardiac interoceptive accuracy in ASD (22), we explored the neural basis of interoception, and the relationship between neural responses and age in a cross-sectional sample of individuals with both typical development and ASD, spanning middle childhood through middle adulthood. We first focused on the insula as the proposed “hub” of interoceptive processing (26–28), and then further expanded our investigation into a whole-brain search, incorporating age and diagnostic status in our model. Given the links between interoception, insula response, and socio-emotional behavior, we predicted that interoceptive-specific response in the insula would be related to ASD symptomatology.

## Methods

### Participants

The Vanderbilt University Medical Center Institutional Review Board approved this study. Informed consent and/or assent was obtained for every participant. In total, 48 individuals with ASD and 60 individuals in a typically developing comparison group (TC) were recruited for this study. Data from 8 participants were not included in the analyzed dataset: 5 participants were excluded for excessive motion (see motion exclusion criteria in Methods, 1 ASD, 4 TC, ages 8-14), and 3 participants had significant acquisition errors during the scan (1 ASD, 2 TC, ages 8-30). The final analyzed sample included 46 individuals with ASD (age 8 to 54, mean= 19.43±10.68 years) and 54 individuals in a typically developing comparison group (TC, age 8 to 53, mean= 21.43±10.41 years). A summary of demographic information is provided in Table 1.

**Table 1.**
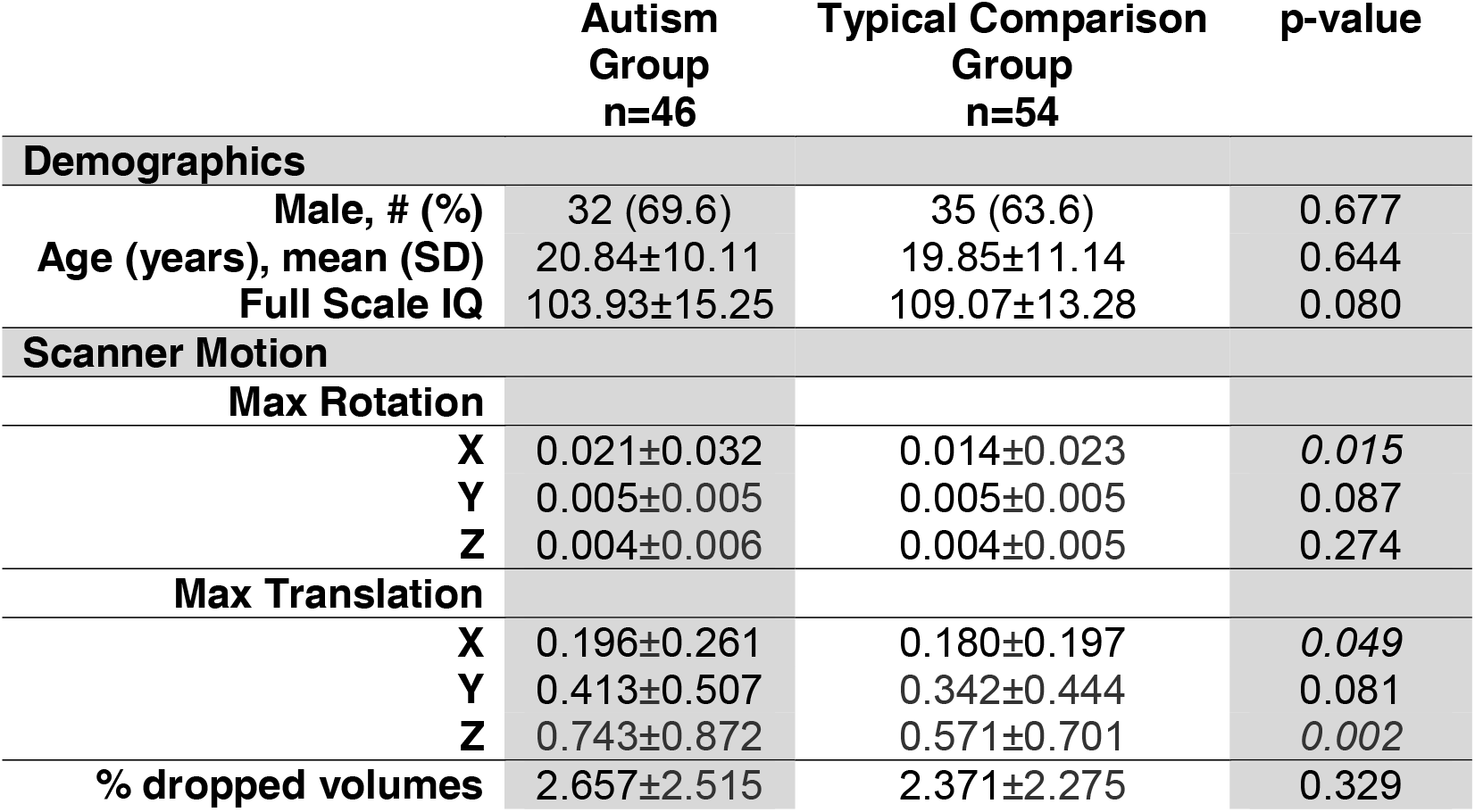
Participant demographic and motion characteristics.

|  | Autism Group<br>n=46 | Typical Comparison Group<br>n=54 | p-value |
| --- | --- | --- | --- |
| **Demographics** |  |  |  |
| Male, # (%) | 32 (69.6) | 35 (63.6) | 0.677 |
| Age (years), mean (SD) | 20.84±10.11 | 19.85±11.14 | 0.644 |
| Full Scale IQ | 103.93±15.25 | 109.07±13.28 | 0.080 |
| **Scanner Motion** |  |  |  |
| **Max Rotation** |  |  |  |
| X | 0.021±0.032 | 0.014±0.023 | 0.015 |
| Y | 0.005±0.005 | 0.005±0.005 | 0.087 |
| Z | 0.004±0.006 | 0.004±0.005 | 0.274 |
| **Max Translation** |  |  |  |
| X | 0.196±0.261 | 0.180±0.197 | 0.049 |
| Y | 0.413±0.507 | 0.342±0.444 | 0.081 |
| Z | 0.743±0.872 | 0.571±0.701 | 0.002 |
| % dropped volumes | 2.657±2.515 | 2.371±2.275 | 0.329 |

All participants had IQ>70, as measured by Wechsler Abbreviated Scales of Intelligence—Second Edition (WASI-II; Wechsler, 2011). ASD diagnosis was confirmed by research-reliable administration of the Autism Diagnostic Observation Schedule— Second Edition (ADOS-2; (32)), and the judgment of a licensed clinical psychologist based on DSM-5 criteria. TC individuals were excluded if they had a history of any psychiatric or learning disorder or had a first-degree relative with ASD. All participants had normal or corrected-to-normal vision. Exclusion criteria for both groups included genetic conditions, neurological disorders, significant head injuries, MRI contraindications, and a number of psychotropic medications (including selective serotonin reuptake inhibitors, serotonin and noradrenergic reuptake inhibitors, noradrenergic and specific serotonergic antidepressants, norepinephrine-dopamine reuptake inhibitors, tricyclics, atypical antipsychotics, mood stabilizers and/or anticonvulsants, anti-hypertensives, and anti-anxiety medications; see Supplemental Table 1 for complete list). Stimulant medications were permitted, but suspended for at least 24 hours prior to the scan. The Social Responsiveness Scale (SRS) (33), a 65-question survey assessing traits relevant for ASD, was administered. For children, a parent or caregiver completed the parent-report SRS version. For adults, either a parent or spouse reported (relative/other report version of SRS, n=9) or participants completed the self-report version. Some individuals did not complete the SRS (total analyzed sample, children: ASD=17, TC=15; adults: ASD=19, TD=16).

### Heartbeat Counting Task

During the functional scan, participants completed a modified cardiac interoceptive accuracy task (7) adapted from (10, 11). The task was presented using e-Prime 2.0 and consisted of 4 runs, with 6-8 randomized blocks of counting heartbeats or a visual stimulus. Instructions were as follows: “*You are going to see either a heart or a star. Each stimulus will be preceeded by a fixation cross. Please stare at the cross until the picture appears. When a heart appears, focus on counting the number of times your heart beats. When a star appears, focus on counting the number of times the star appears.*” During the heartbeat condition, a heart shape was static on the screen and individuals were instructed to count their own heartbeat while lying still in the scanner. During the visual condition, a low-contrast star shape flashed at a rate similar to an average resting heart rate. The flashing visual counting stimulus was designed to be difficult to detect visually. Following each block, participants reported their count using hand signals to a researcher in the scanning room. There was a 15 second rest between each block. This task was practiced outside of the scanner to ensure task comprehension and participants were continually reminded that they were not to physically feel any part of their body for a pulse.

During the entire scan, the participant’s heart rate was recorded using the peripheral pulse oximeter associated with the 3.0 Tesla Phillips Achieva MRI scanner (sampling rate of 0.002 seconds). Accuracy on the heartbeat task was calculated by comparing the participant counts to the physiological heart rates from the scanner. Accuracy was calculated on each trial, or block during each scanner run as the absolute value of ((actual heartbeats from scanner output - reported heartbeats) / actual heartbeats). Overall accuracy was based on the average for all completed blocks of heartbeat counting. Accuracy on the visual counting task was calculated as follows: absolute value of ((actual flashes in e-prime output – reported flashes) / actual flashes in e-prime output). Overall accuracy was based on the average for all completed blocks of visual counting. Some visual (n=9) and heartbeat (n=18) counting data were missing due to technical issues during data transfer from the scanner. Physiological recording data were cleaned prior to analysis; recordings of fewer than 20 beats per 30 seconds were rejected as unlikely to reflect accurate heartrates. Similarly, participant-report counts that were below 5 were rejected as inconsistent with task understanding. Counts were not used if the participant indicated they were falling asleep or unable to sustain attention for a particular block. For these reasons, 5 participants’ accuracy data were not included in the final analysis. The final sample of accuracy data included for heartbeat accuracy: ASD, n=28, TC, n=41 and for visual accuracy: ASD, n=29, TC, n=44.

### Image Acquisition

Image data was collected using a 3.0 Tesla Phillips Achieva MRI scanner with a 32-channel SENSE head coil. A high-resolution T1-weighted anatomical image was acquired (1mm^3^ resolution, TR=8 msec, TE= 3.7 msec, acquisition matrix=256×256) for registration of functional images. Functional images were acquired with a whole-brain T2*-weighted EPI sequence (axial slices, voxel size=2.5×2.5×3mm, TR=2 s, TE=25 msec, FOV= 96×96, flip angle=90°).

### Image Preprocessing and Analysis

For processing anatomical images, we used a large-scale neuroimaging data platform (34). A multi-atlas segmentation algorithm (35) with a set of manually labeled atlases (Neuromorphometrics, Inc., Somerville, MA, USA) was applied to the T1-weighted image in order to obtain a gray matter image. All gray matter regions were summed to make a total gray matter image, and then the gray matter image was nonlinearly normalized to a MNI-152 gray matter probabilistic template (OASIS project (http://www.oasis-brains.org), provided by Neuromorphometrics, Inc. (http://www.neuromorphometrics.com)). The registration algorithm minimized the sum of squared differences between gray matter image and template (SPM12 “Old Normalise”, (36, 37)) and modeled spatial nonlinearity with a discrete cosine basis of 25 mm lower cutoff. Using parameters resulting from the anatomical normalization step, each subject’s coregistered functional images were then warped to MNI space.

FMRI data processing was carried out using FEAT (FMRI Expert Analysis Tool) Version 6.00, part of FSL (FMRIB’s Software Library, www.fmrib.ox.ac.uk/fsl). Motion correction was conducted with MCFLIRT (38); brain extraction with BET (39), a high-pass filter (300s), grand-mean intensity normalization of the entire 4D dataset by a single multiplicative factor and spatial smoothing (FWHM=5mm). Time-series statistical analysis was conducted with prewhitening using FILM with local autocorrelation correction (40).

Each block was modeled as a single contrast of stimulus – baseline (rest of 15s), convolved with a double-gamma hemodynamic response function with temporal filtering added to the GLM model. Standard motion parameters were included in the general linear model (GLM), with the addition of DVARS (D, temporal derivatives of time courses; VARS, variance of root mean squares of head motion across voxels) and framewise displacement metrics for head motion (41) to the GLM as confound explanatory variables (EV) to remove effects of outlier volumes from the parameter estimates of interest. Individual runs were rejected based on peak motion > 5mm or if DVARS/FD metrics flagged more than 10% of volumes as outliers. In the analyzed dataset, there were no differences in the number of outlier volumes by diagnostic group. There was a significant difference in motion by diagnostic group such that ASD had higher max translation in x (p=0.049) and z (p=0.002) and rotation around x (pitch, p=0.015).

Second-level analyses combined runs for each subject using a fixed-effects model, by forcing random effects variance to zero in FLAME (FMRIB’s Local Analysis of Mixed Effects) (42–44).

### Region of Interest Analysis

Regions of interest (ROI) in this study included sub-divisions of the bilateral insula as defined cyto-architecturally (45), as previous work has suggested different interoceptive functions per laminar architecture (46). The Farb et al. insular ROIs include 16 subdivisions on right (8) and left (8) insular cortex. Percent signal change was extracted using featquery in FSL for each of the 16 insular ROIs and then averaged into 3 ROIs on each insular cortex: anterior (3 most anterior ROIs), mid (2 middle ROIs), and posterior (3 most posterior ROIs). Figure 1 shows a depiction of these bilateral insular subdivisions.

**Figure 1.**
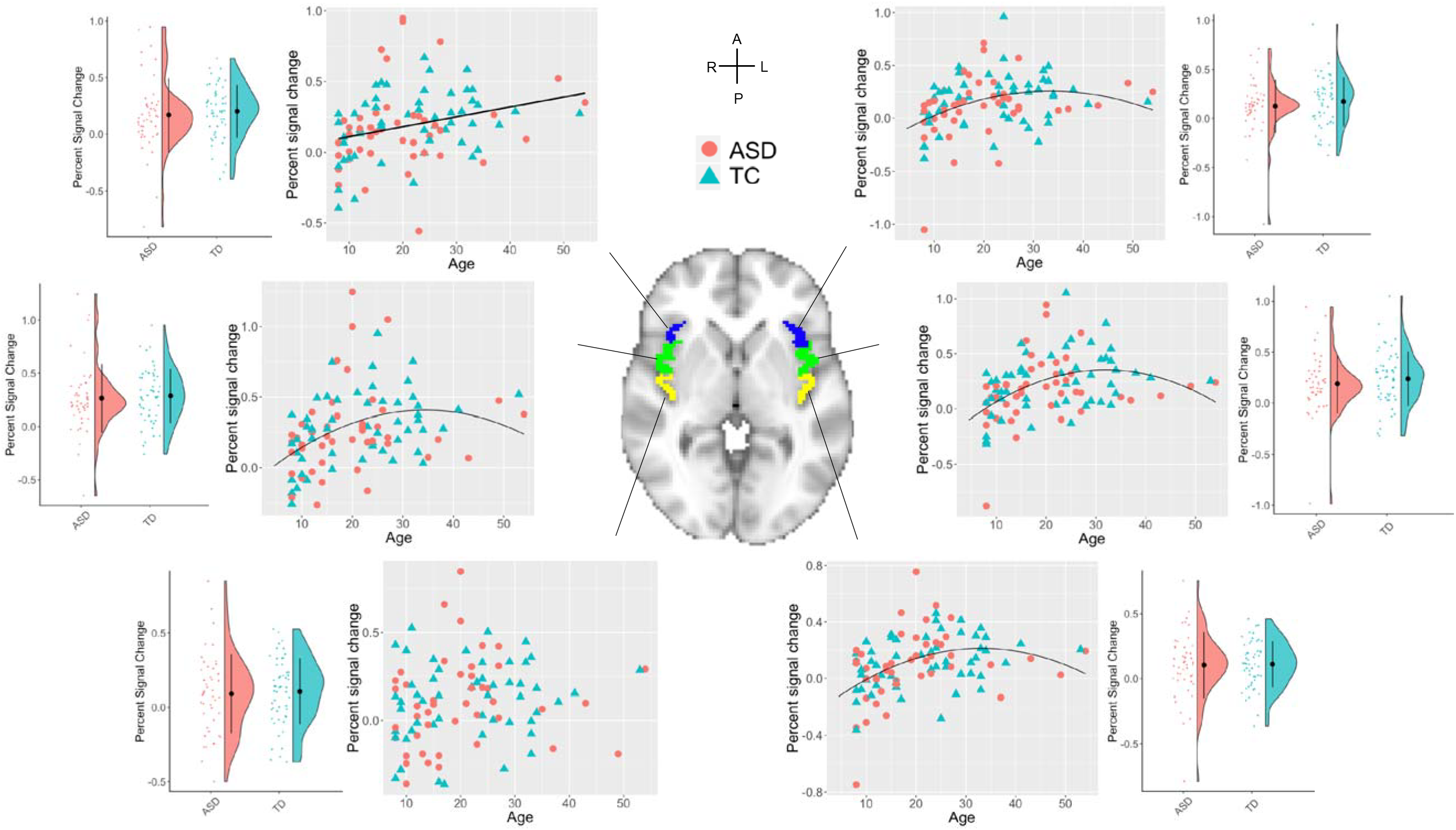
Percent signal change in subdivisions of the bilateral insula (heartbeat counting-visual counting) is not different by diagnostic group but is related to age. Regions of interest are shown in the middle on the MNI template brain. Insula regions are subdivided into 3 regions on each side, blue: anterior, green: mid, yellow: posterior insula. **There were no differences in percent signal change in any of the insula subregions by diagnostic group:** for anterior, left: ASD=0.124±0.325, TD=0.171±0.233, W=1086, p=0.353, and right: ASD=0.169±0.326, TD=0.202±0.233, W=1052, p=0.243; for middle, left: ASD=0.190±0.291, TD=0.238±0.265, W=1068, p=0.291, and right: ASD=0.265±0.321, TD=0.287±0.253, W=1099, p=0.402; and posterior, left: ASD=0.105±0.254, TD=0.111±0.177, W=1211, p=0.958, and right: ASD=0.091±0.266, TD=0.107±0.222, W=1139, p=0.577. **For relationships with age**, model fit and parameters for each insula subdivision can be found in Table 2.

### Whole-brain Group Analysis

Group analyses were carried out using FLAME (FMRIB’s Local Analysis of Mixed Effects) stage 1 model (42–44). Given the developmental age range in this sample, age was considered a covariate in the GLM, as both a linear and nonlinear (quadratic) association. All group maps were made using a cluster threshold of z = 2.3 and a p-value = 0.05 (family-wise error corrected based on clusters defined by Gaussian random field theory (47)). Additionally, group contrasts of ASD>TD and TD>ASD were examined. Nonparametric permutation testing of group results was conducted using FSL’s randomize (48). Using the Threshold-Free Cluster Enhancement method, 5000 iterations of each statistics map were generated, results are reported at the *p*<0.05 level.

### Statistical Analysis

Demographic and survery data for this study was collected and managed with REDCap electronic data capture tools (49). Data processing and statistical analyses were conducted using Python (v.2.7) and R (v.3.3.2). Multiple comparison corrections were calculated using the Holm method (50).

To examine diagnostic group differences in demographic variables, motion parameters, and percent signal change in insula subdivisions, we used parametric tests (t-tests), non-parametric tests (Mann-Whitney), or chi-square tests, where appropriate. To investigate associations between age and percent signal change in insula subdivisions, we first investigated nonlinear, quadratic fit regression models and then calculated spearman’s rho for those insula subdivisions where a nonlinear, quadratic fit was not significant.

To address missing data in interoceptive accuracy data and SRS scores, we used multiple imputation by chained equations (51, 52); data were imputed to 5 datasets, with up to 40 iterations, using predictive mean matching and estimates from all reported tests were pooled. To investigate relationships between heartbeat counting accuracy, age, and IQ, we calculated spearman’s rho. To compare group means in heartbeat counting accuracy, we calculated student’s t-tests.

To investigate effects of neural responses in the insula, diagnostic group, and age on social symptomology, we employed linear regression models with SRS total T-scores as the dependent variable. Age, diagnostic group, and percent signal change from the left posterior insula were entered into the model, as well as a diagnostic group by insula variable. Models were calculated separately in self- and other-reporter versions of the SRS.

## Results

### Cardiac interoceptive accuracy

Heartbeat counting accuracy did not differ by group (ASD: 71.50 ± 26.59%, TC: 75.71 ± 20.09%, t=−0.936, df=51.54, p=0.354), and reached accuracy levels consistent with our previous results outside of the scanner (10). Age was not significantly associated with heartbeat accuracy in the total sample (r=0.114, 95% CI (−0.140-0.355), p=0.379); additionally, there was no significant interaction between age and diagnostic group on heartbeat accuracy (age*diagnosis: ß=−0.021, p=0.957). Heartbeat counting accuracy did not differ by gender (Female: 74.09 ± 24.42%, Male: 73.62 ± 22.09%, t=0.102, df=40.56, p=0.919). IQ was significantly correlated with heartbeat accuracy across both groups (r=0.243, 95% CI (0.024-0.439), p=0.030). Using linear regression, we investigated the possibility of an interaction between IQ and diagnostic group on heartbeat accuracy, correcting for age (r^2^=0.122). IQ was still associated with heartbeat accuracy (ß=0.502, p=0.014); age (ß=0.226, p=0.283), diagnostic group (ß=39.52, p=0.180), and diagnostic group*IQ (ß=−0.351, p=0.205) were not significant predictors of heartbeat accuracy.

In the control condition, mean visual counting accuracy was over 92% for both groups with no significant group differences (ASD: 92.61 ± 16.47%, TC: 93.60 ± 13.90%, t=−0.329, df=43.80, p=0.744). Visual counting accuracy was correlated with heartbeat counting accuracy (r=0.286, p=0.026), however, when visual counting accuracy and IQ were in the same linear regression model, IQ was still associated with heartbeat counting accuracy (ß=0.314, p=0.046) while visual counting accuracy was not (ß=0.121, p=0.552, r^2^=0.083).

### Insula response during interoception

We examined percent signal change in subdivisions of the insula as it related to diagnosis and age. For each subdivision of the insula, mean percent signal change was not significantly different by diagnostic group, for anterior, left: ASD=0.124±0.325, TD=0.171±0.233, W=1086, p=0.353, and right: ASD=0.169±0.326, TD=0.202±0.233, W=1052, p=0.243; for middle, left: ASD=0.190±0.291, TD=0.238±0.265, W=1068, p=0.291, and right: ASD=0.265±0.321, TD=0.287±0.253, W=1099, p=0.402; and posterior, left: ASD=0.105±0.254, TD=0.111±0.177, W=1211, p=0.958, and right: ASD=0.091±0.266, TD=0.107±0.222, W=1139, p=0.577 (Figure 1).

For age, we explored both linear and nonlinear, quadratic models as previous studies have demonstrated both linear and nonlinear developmental trajectories in the brain (53). Figure 1 shows the inverted curvilinear relationship between age and percent signal change in 4 of the 6 insular subdivisions. Of the remaining 2 subdivisons, the right anterior insula had a better fit with a linear model, whereas the right posterior insula did not show a significant relationship with age in either linear or nonlinear models. Table 2 shows the best-fit model parameters for each ROI and age.

**Table 2.**
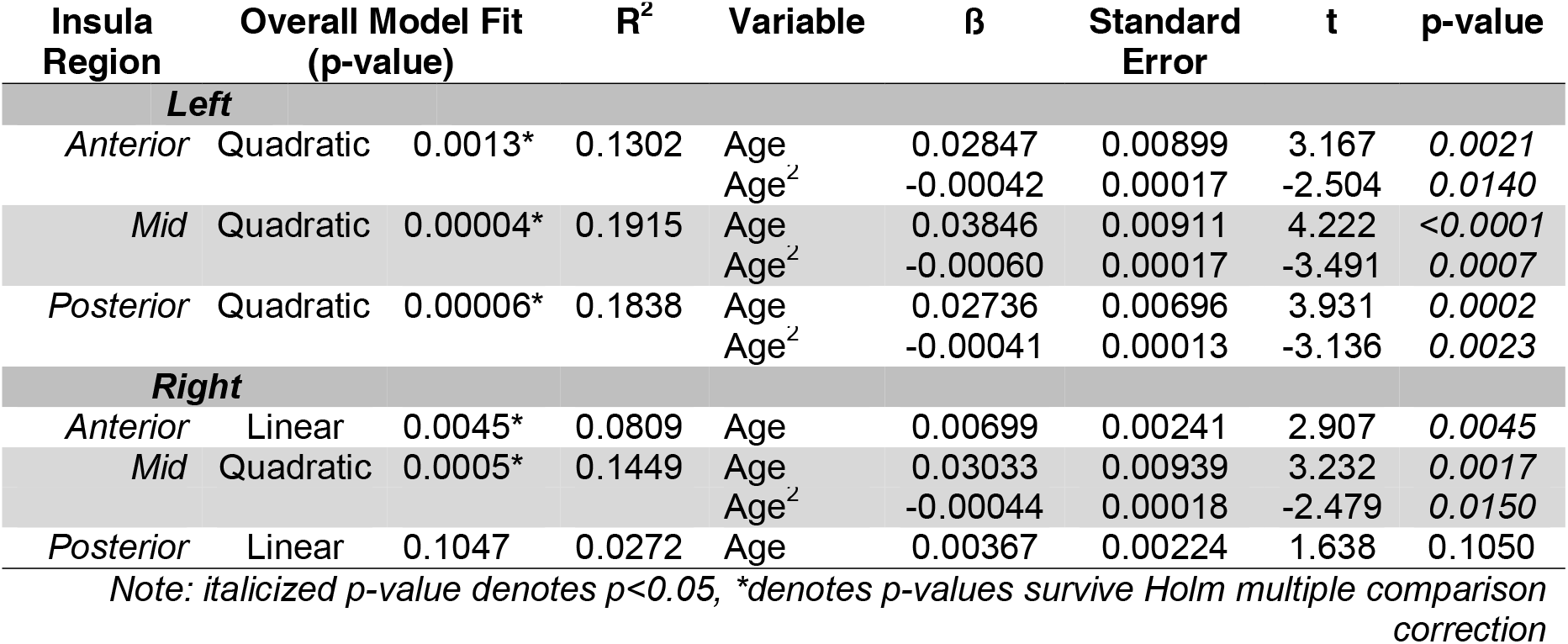
Linear and nonlinear models of age by percent signal change in insular regions.

There were no significant relationships between percent signal change in insula subregions and heartbeat counting accuracy ([anterior: left, r=−0.020,p=0.865; right, r=0.132, p=0.268]; [mid: left, r=0.046, p=0.692; right, r=0.105, p=0.372]; [posterior: left, r=0.079, p=0.484, right: r=0.006, p=0.958]).

### Interoception-specific whole brain responses

For the ASD and TC groups, interoception-specific BOLD response was observed bilaterally in the insula, medial frontal gyrus, anterior cingulate, primary and secondary somatosensory regions, visual cortex, and cerebellum (Figure 2). We also investigated group differeneces: there were no significant clusters where ASD>TC or TC>ASD. There were several responses associated with age, both with a positive linear association (bilateral anterior, mid, and posterior insula, secondary somatosensory, anterior cingulate) as well as with a negative quadratic association (left insula). Following permutation testing, both groups retained clusters in the bilateral insula, while the ASD group also showed response in the visual cortex and cerebellum. In the permutation testing, positive linear associations with age expanded to include the majority of the interoceptive network (Supplemental Figure 1). There were no clusters related to quadratic, nonlinear age effects.

**Figure 2.**
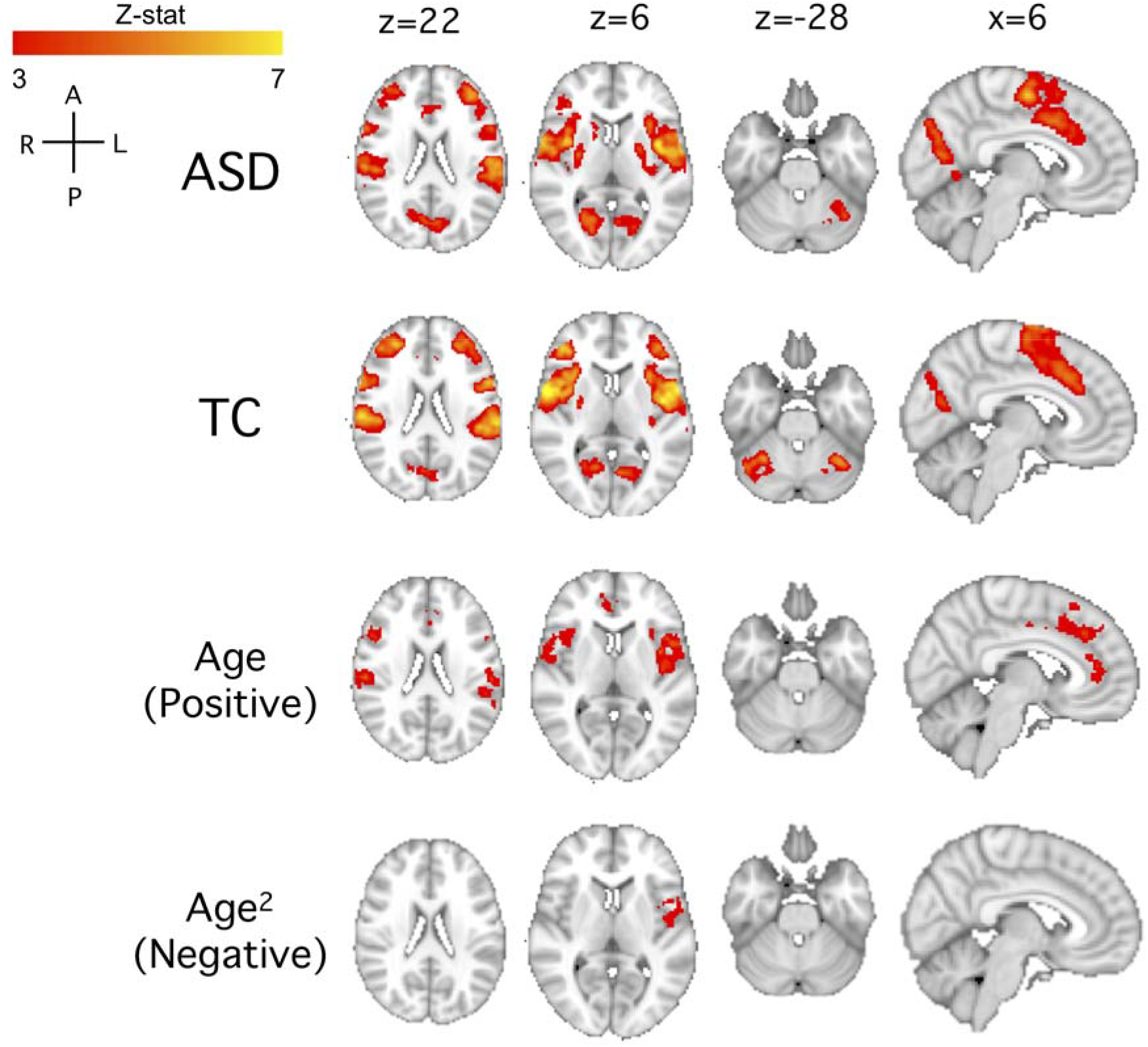
Whole brain heartbeat counting-visual counting in the total sample. Group maps are shown for the autism spectrum disorders (ASD), typical comparison group (TC), and linear and nonlinear age associations. Group maps were calculated with z>3 and p<0.01. We also tested group comparison of ASD>TC and TC>ASD (thresholded at z>2.3 and p<0.05), but there were no significant clusters. There were no negative linear age associations and no positive nonlinear age^2^ associations. See Supplemental Figure 1 for permutation testing results.

### Associations of interoceptive response and autism-related behaviors

We investigated the relationship between insular response during our interoception task and ASD-related symptomology, using SRS total T-score as the dependent variable in separate linear regression models by self and other report. We chose the left postserior insula for this analysis, because posterior insula contains primary interoceptive cortex and left insula may be more responsive to social and other emotionally relevant stimuli (54, 55). As age was significantly associated with posterior insula response, we included age in our models of SRS scores. We also incorporated an interaction between insula response and diagnostic group in our models.

Table 3 summarizes models for total SRS scores, separated by self-report and other reporters. In the self-report model, there was a significant interaction between percent signal change in the left posterior insula and diagnostic group on total SRS T-scores (Figure 3, Table 3). There was a positive relationship between left posterior insula response and SRS scores for those with ASD, whereas there was a negative relationship between insula response and SRS scores in the TC group. In the other-reporting model, diagnostic group was significantly associated with SRS scores, but left posterior insula response was not significantly related, nor was there a significant interaction between left posterior insula response and diagnostic group. Age was not significantly associated with SRS score in either model.

**Table 3.** Linear regression models of ASD-related social symptomology with left posterior insula and diagnostic group.

| $R^2$ | Variable | $\beta$ | Standard Error | t | p-value |
| --- | --- | --- | --- | --- | --- |
| ***Self-Report*** |  |  |  |  |  |
| 0.801 | Age | 0.1224 | 0.307 | 0.399 | 0.6947 |
|  | Left Posterior Insula (% Signal Change) | 41.407 | 13.356 | 3.100 | *0.0058* |
|  | Diagnostic Group | -16.363 | 4.461 | -3.668 | *0.0016* |
|  | Insula x Group | -65.450 | 17.023 | -3.844 | *0.0011* |
| ***Other-Report*** |  |  |  |  |  |
| 0.591 | Age | -0.053 | 0.202 | -0.264 | 0.7926 |
|  | Left Posterior Insula (% Signal Change) | -4.253 | 7.539 | -0.564 | 0.5754 |
|  | Diagnostic Group | -21.840 | 2.937 | -7.435 | *<0.0001* |
|  | Insula x Group | -1.021 | 13.310 | 0.077 | 0.9392 |

*Note: italicized p-value denotes $p < 0.05$*

**Figure 3.**
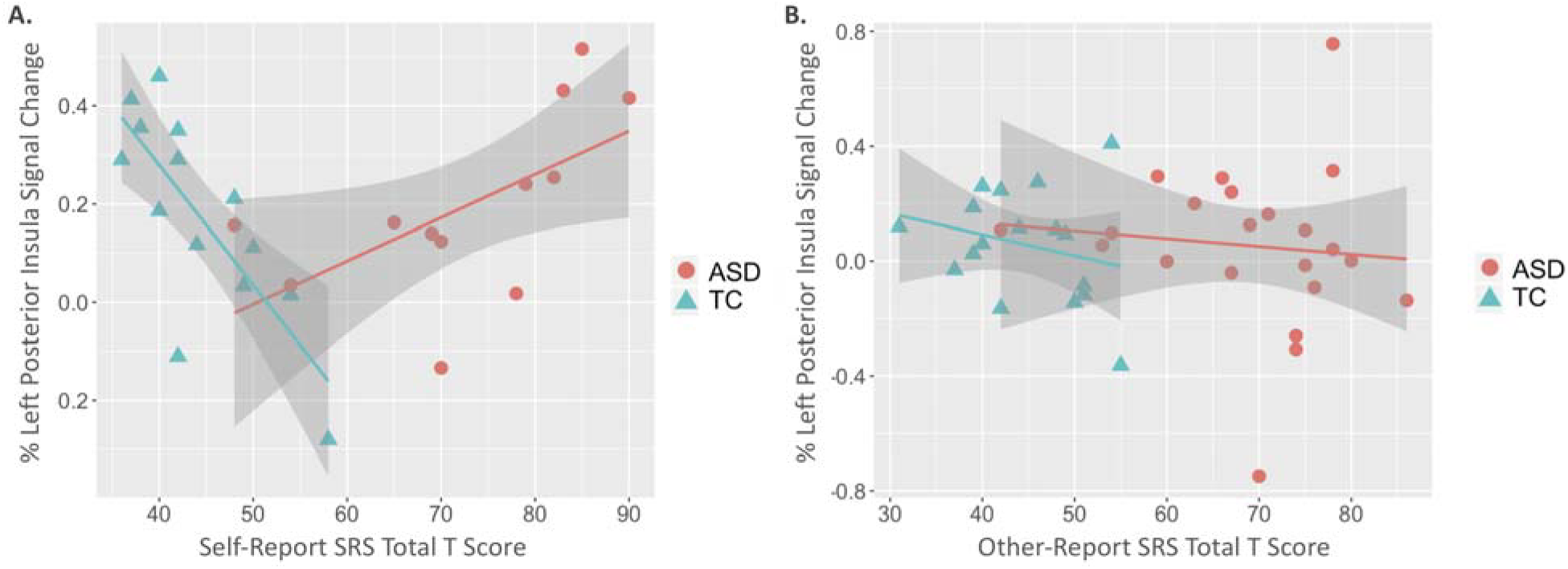
Associations between autism-relevant symptomology and left posterior insular response during an interoceptive task. In the self-report model (A), there was a significant interaction between percent signal change in the left posterior insula and diagnostic group on total SRS T-scores. In the other-reporting model (B), diagnostic group was significantly associated with SRS scores, but left posterior insula response was not significantly related, nor was there a significant interaction between left posterior insula response and diagnostic group. See Table 3 for model specifics.

## Discussion

In this study, we report developmental effects on neural response to an interoceptive heartbeat counting task in children and adults with and without ASD. Neural responses throughout the insula during the heartbeat counting task were not different by diagnostic group, consistent with our previous behavioral findings of intact cardiac interoception in ASD (Schauder et al., 2015, Mash et al., 2017). However, we found that the response of primary interoceptive cortex during interoception interacted with group to predict ASD symptoms as measured by SRS self-report. These findings suggest that people with autism may integrate internal sensory cues and external, social information, in a different way. This work reinforces the link between neural processing of internal sensory cues and social-emotional awareness. The size and broad age range of our sample provided a unique opportunity to detect age-related patterns in interoception.

In this study, there was a significant relationship between interoceptive processing and age that was different for children compared to adults. There was a positive relationship between age and insular response to heartbeat counting that continued into early adulthood. This mirrors the findings of Li and colleagues, (24), who described nonlinearly increasing interoceptive BOLD responses in the ventral anterior insula with age in a sample aged 10-20 years. Our results corroborate this peak of insular response in late adolescence or early adulthood, which could suggest maturation of these circuits in this timeframe. Interestingly, cardiovagal autonomic function peaks around the same time, suggesting maturation of important autonomic neural mechanisms in parallel (56). While our sample did include up to middle adulthood, it is not clear how this relationship might extend into later adulthood. There is evidence of reduced interoceptive accuracy with age in adults (57, 58), but it is not clear how, or if, the neural circuits underlying interoception change throughout adulthood. Our results suggest there is a tempered decline in interoceptive responses in the insula, but replication is needed to confirm this finding.

There were no significant differences in cardiac interoceptive neural processing in ASD, either in insula subdivisions or in the whole brain search, and no group differences in interoceptive accuracy. Yet, we show a clear relationship between neural processing of interoceptive signals and reported social responsiveness in ASD. While some previous studies have reported diminished heartbeat counting accuracy in children (16) and adults with ASD (17, 18), we found no group differences in the current study, which is consistent with the majority of reports (10, 11, 19–21). These data may suggest that, globally, primary cardiac interoceptive ability is not dramatically impacted in ASD. However, several previous reports suggest differences in interoceptive sensitivity and awareness, rather than accuracy, reflecting altered attention to and interpretation of interoceptive cues (9, 17, 18, 59). This mixed profile, potentially reflecting a mismatch between accuracy and confidence (60) could significantly impact social-emotional awareness and development by increasing the likelihood of erroneous or inconsistent interoceptive interpetations. This is supported by our data in adults, where insula response was related to self-reported social functioning. Yet, we saw no relationship in caregiver reports of social functioning. Its important to note that self-report was only collected in adults and the majority of caregiver reports were collected in children and young adults. More research is needed to understand the direct influence of primary interoceptive insula response on interoceptive sensitivity and awareness, and how that influence changes during development.

In our study, there were no differences in accuracy by diagnostic group, gender, or age. However, accuracy was significantly related to IQ. This may reflect a general understanding of task requirements, but more than likely reflects participants’ prior knowledge of average resting heartrate (see (61, 66) for more). As we did not measure beliefs regarding average heartrate, the role prior knowledge may play is unclear. There is likely a complicated association between interoceptive accuracy, age (58), IQ (61), and diagnostic group, as demonstrated in our previous work (10, 11), but our study was likely not powered to detect these relationships. To our knowledge, no other studies have examined associations between insula percent signal change and cardiac interoceptive accuracy.

There are important strengths and limitations to consider in this study. This is one of a few studies to experimentally measure interoceptive accuracy and neural correlates in children, which adds to the novelty of this study. Additionally, we excluded for a number of medications that could influence autonomic and/or higher-order cognitive or affective function. In a sample with ASD, this is very difficult given the common use of these types of medications. Yet, because of the common use of these medications, future studies will need to understand the interoceptive consequences of these medications on socio-emotional health. As this work is cross-sectional, longitudinal data are needed to confirm developmental shifts in interoception-related BOLD response. We also only investigated one modality of interoception, however, there are a number of other internal stimuli highly related to development and ASD, like gastrointestinal-related cues(62) and pain (63, 64). There is limited evidence that interoceptive accuracy in one modality like heartbeat is related to accuracy in other modalities (65), yet this has not been investigated developmently or in ASD specifically. Similarly, heartbeat counting accuracy may not be as ‘purely’ interoceptive as it appears; it may be influenced by non-interoceptive processes such as beliefs about heart rates, estimating heart beats, etc. (see (66–69)). However, for this study, this task provided a developmentally accessible interoceptive task to examine neural correlates of interoceptive processes without the additional consideration of exteroceptive signals as in heartbeat discriminiation tasks. Given wide-ranging implications of interoceptive processing across development and in ASD, it will be important to address these limitations in future studies.

Another important consideration when interpreting interoceptive processing in socio-emotional functioning is the role of alexythmia. Alexithymia refers to “impaired awareness of emotions due to a deficit in processing of affective information” (70) and is highly prevalent in ASD (70). In fact, Bird and colleagues suggest interoceptive deficits are only observable in the subset of people with ASD who are alexithymic (19, 20, 71). However, recent work suggests that cardiac interoceptive accuracy is not associated with ASD traits or alexithymia (21). While we did not measure alexithymia, we did not find differences in cardiac interoceptive accuracy by diagnostic group. The overlapping features of ASD and alexithymia clearly complicate this conceptual landscape and likely contribute to mixed findings.

Given our current data, we suggest that 1) there is a developmental trajectory of interoceptive neural processing in the insular cortex that is dynamic throughout early adulthood, and 2) interoceptive influence on social functioning in ASD is not likely due to primary interoceptive deficits. Future studies will need to examine connectivity of interoceptive networks, especially in the face of exteroceptive demands, to understand influences on social behaviors. Given evidence for altered structural connectivity in the insula of children with ASD (12), this will be an important consideration moving forward. Greater relative involvement of extrastriate visual cortex in individuals with ASD across the lifespan may further suggest differences in strategies for bodily perception that could be explored in the context of internalizing, anxiety, and other mental health variables. Thus, future studies should examine how people access interoceptive memories or information (72), as this may be highly relevant in social functioning. Finally, longitudinal studies of interoception will help to shed light on the complex interplay between visceral sensation, mental health, and social function as these processes evolve over the lifespan.

## Acknowledgements

We also acknowledge the support from the Vanderbilt Kennedy Center Treatment and Research Institute for Autism Spectrum Disorders (NICHD U54HD083211) and Vanderbilt Institute for Clinical and Translational Research grant support (UL1 TR000445 from NCATS/NIH). We thank Norman Farb for sharing regions of interest in the insula and for his guidance in their implementation.

---

**Supplemental Figure 1.**
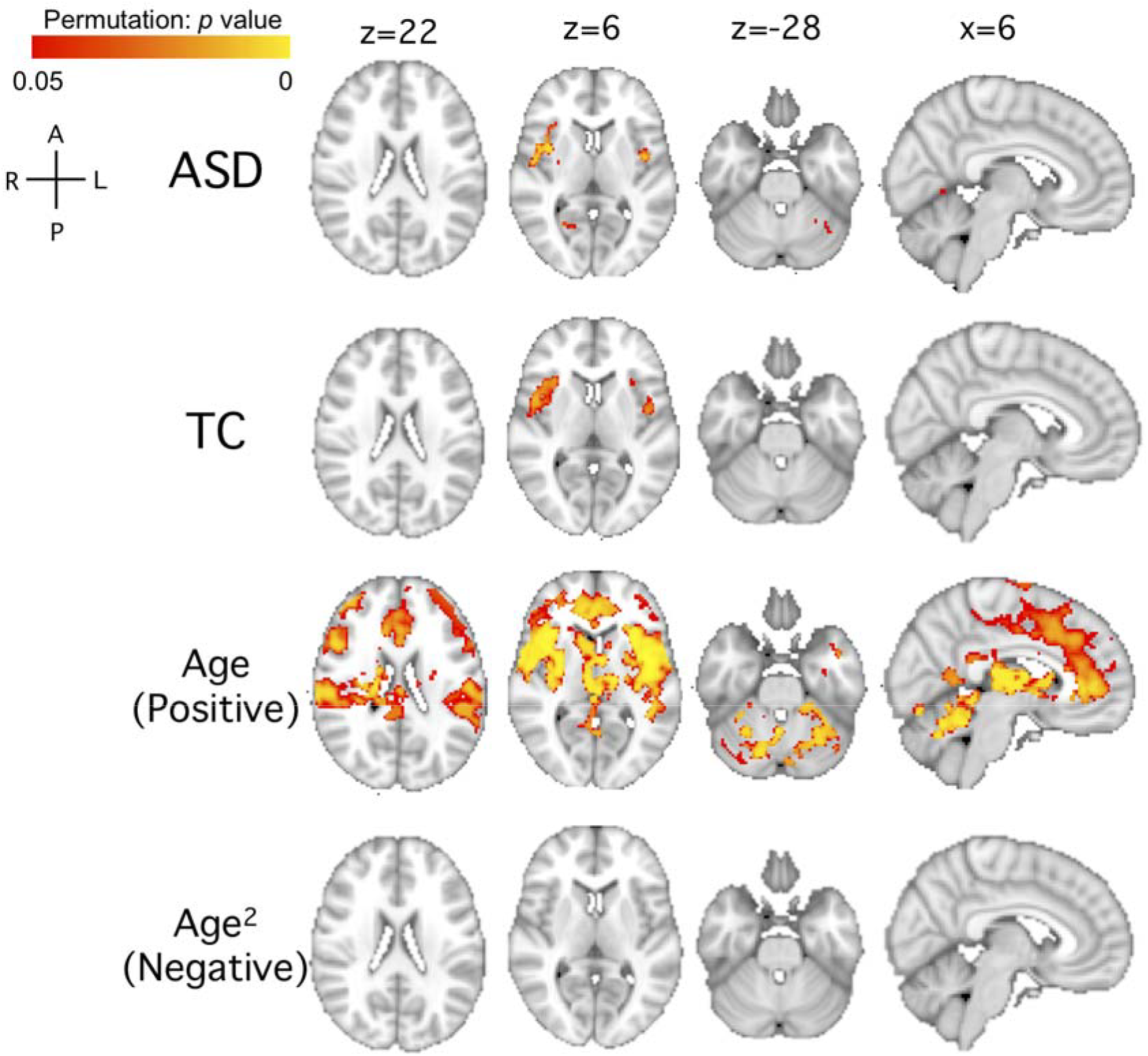
Whole brain heartbeat counting-visual counting in the total sample permutation tested. Group maps are shown for the autism spectrum disorders (ASD) group, typical comparison (TC) group, and linear and nonlinear (quadratic) age associations. Group maps were calculated with z>3 and p<0.01. We also tested group comparison of ASD>TC and TC>ASD (thresholded at z>2.3 and p<0.05) but there were no significant clusters. There were no negative linear age associations and no positive nonlinear age^2^ associations. Nonlinear permutation tested clusters were calculated using fsl randomise. Reported clusters represent p<0.05 after 5000 iterations.

**Supplemental Table 1.**

| **Excluded Psychotropic Medications** |  |  |  |
| --- | --- | --- | --- |
| **Antidepressants** | SSRI | Prozac | fluoxetine |
|  |  | Zoloft | sertraline |
|  |  | Luvox | fluvoxamine |
|  |  | Paxil | paroxetine |
|  |  | Celexa | citalopram |
|  |  | Lexapro | escitalopram |
|  | SNRI | Effexor | venlafaxine |
|  |  | Cymbalta | duloxetine |
|  | NASSA | Remeron | mirtrazapine |
|  | NADRI | Wellbutrin | bupropion |
|  | Tricyclics | Elavil | amitriptyline |
|  |  | Tofranil | imipramine |
|  |  | Anafranil | clomipramine |
| **Other** | atypical antipsychotics | Risperdal | risperdone |
|  |  | Seroquel | quetiapine |
|  |  | Abilify | aripiprazole |
|  |  | Geodon | ziprasidone |
|  |  | Zyprexa | olanzapine |
|  | mood stabilizers and/or anticonvulsants | Lithium | lithium carbonate |
|  |  | Tegretol | carbamazepine |
|  |  | Trileptal | oxcarbazepine |
|  |  | Depakote | divalproex sodium |
|  |  | Lamictal | lamotrigine |
|  |  | Neurontin | gabapentin |
|  |  | Topamax | topiramate |
|  |  | Keppra | levetiracetam |
|  | anti-hypertensive | Clonidine | clonidine |
|  |  | Catapres | clonidine |
|  |  | Intuniv | guanfacine |
|  |  | Tenex | guanfacine |
|  | anti-anxiety | Xanax | alprazolam |
|  |  | Klonopin | clonazepam |
|  |  | Ativan | lorazepam |
|  |  | Valium | diazepam |

Note: SSRI, selective serotonin reuptake inhibitors; SNRI, serotonin and noradrenergic reuptake inhibitors; NASSA, noradrenergic and specific serotonergic antidepressants; NADRI, norepinephrine-dopamine reuptake inhibitors.

## References

1. Craig AD (2002): How do you feel? Interoception: the sense of the physiological condition of the body. Nat Rev Neurosci. 3: 655–666.

2. Pollatos O, Schandry R (2008): Emotional processing and emotional memory are modulated by interoceptive awareness. Cogn Emot. 22: 272–287.

3. Critchley HD, Garfinkel SN (2017): Interoception and emotion. Curr Opin Psychol. 17: 7–14.

4. Palmer CE, Tsakiris M (2018): Going at the heart of social cognition: is there a role for interoception in self-other distinction? Curr Opin Psychol. 24: 21–26.

5. Murphy J, Brewer R, Catmur C, Bird G (2017): Interoception and psychopathology: A developmental neuroscience perspective. Dev Cogn Neurosci. 23: 45–56.

6. Maister L, Tang T, Tsakiris M (2017): Neurobehavioral evidence of interoceptive sensitivity in early infancy. eLife. 6. doi: 10.7554/eLife.25318.

7. Schandry R (1981): Heart Beat Perception and Emotional Experience. Psychophysiology. 18: 483–488.

8. Koch A, Pollatos O (2014): Cardiac sensitivity in children: sex differences and its relationship to parameters of emotional processing. Psychophysiology. 51: 932–941.

9. Fiene L, Brownlow C (2015): Investigating interoception and body awareness in adults with and without autism spectrum disorder. Autism Res Off J Int Soc Autism Res. 8: 709–716.

10. Mash LE, Schauder KB, Cochran C, Park S, Cascio CJ (2017): Associations Between Interoceptive Cognition and Age in Autism Spectrum Disorder and Typical Development. J Cogn Educ Psychol JCEP. 16: 23–37.

11. Schauder KB, Mash LE, Bryant LK, Cascio CJ (2015): Interoceptive ability and body awareness in autism spectrum disorder. J Exp Child Psychol. 131: 193–200.

12. Failla MD, Peters BR, Karbasforoushan H, Foss-Feig JH, Schauder KB, Heflin BH, Cascio CJ (2017): Intrainsular connectivity and somatosensory responsiveness in young children with ASD. Mol Autism. 8: 25.

13. Uddin LQ, Menon V (2009): The anterior insula in autism: under-connected and under-examined. Neurosci Biobehav Rev. 33: 1198–1203.

14. Ondobaka S, Kilner J, Friston K (2017): The role of interoceptive inference in theory of mind. Brain Cogn. 112: 64–68.

15. Quattrocki E, Friston K (2014): Autism, oxytocin and interoception. Neurosci Biobehav Rev. 47: 410–430.

16. Palser ER, Fotopoulou A, Pellicano E, Kilner JM (2018): The link between interoceptive processing and anxiety in children diagnosed with autism spectrum disorder: Extending adult findings into a developmental sample. Biol Psychol. 136: 13–21.

17. Garfinkel SN, Tiley C, O’Keeffe S, Harrison NA, Seth AK, Critchley HD (2016): Discrepancies between dimensions of interoception in autism: Implications for emotion and anxiety. Biol Psychol. 114: 117–126.

18. Mul C-L, Stagg SD, Herbelin B, Aspell JE (2018): The Feeling of Me Feeling for You: Interoception, Alexithymia and Empathy in Autism. J Autism Dev Disord. 48: 2953–2967.

19. Shah P, Hall R, Catmur C, Bird G (2016): Alexithymia, not autism, is associated with impaired interoception. Cortex J Devoted Study Nerv Syst Behav. 81: 215–220.

20. Shah P, Catmur C, Bird G (2016): Emotional decision-making in autism spectrum disorder: the roles of interoception and alexithymia. Mol Autism. 7: 43.

21. Nicholson TM, Williams DM, Grainger C, Christensen JF, Calvo-Merino B, Gaigg SB (2018): Interoceptive impairments do not lie at the heart of autism or alexithymia. J Abnorm Psychol. 127: 612–622.

22. DuBois D, Ameis SH, Lai M-C, Casanova MF, Desarkar P (2016): Interoception in Autism Spectrum Disorder: A review. Int J Dev Neurosci Off J Int Soc Dev Neurosci. 52: 104–111.

23. Crone EA, Dahl RE (2012): Understanding adolescence as a period of social–affective engagement and goal flexibility. Nat Rev Neurosci. 13: 636–650.

24. Li D, Zucker NL, Kragel PA, Covington VE, LaBar KS (2017): Adolescent development of insula-dependent interoceptive regulation. Dev Sci. 20. doi: 10.1111/desc.12438.

25. Pollatos O, Matthias E, Keller J (2015): When interoception helps to overcome negative feelings caused by social exclusion. Front Psychol. 6. doi: 10.3389/fpsyg.2015.00786.

26. Schulz SM (2016): Neural correlates of heart-focused interoception: a functional magnetic resonance imaging meta-analysis. Philos Trans R Soc B Biol Sci. 371. doi: 10.1098/rstb.2016.0018.

27. Craig AD (Bud) (2014): Topographically Organized Projection to Posterior Insular Cortex from the Posterior Portion of the Ventral Medial Nucleus (VMpo) in the Long-tailed Macaque Monkey. J Comp Neurol. 522: 36–63.

28. Gu X, Hof PR, Friston KJ, Fan J (2013): Anterior Insular Cortex and Emotional Awareness. J Comp Neurol. 521: 3371–3388.

29. Garfinkel SN, Critchley HD (2013): Interoception, emotion and brain: new insights link internal physiology to social behaviour. Commentary on:“Anterior insular cortex mediates bodily sensibility and social anxiety” by Terasawa et al. (2012). Soc Cogn Affect Neurosci. 8: 231–234.

30. Terasawa Y, Shibata M, Moriguchi Y, Umeda S (2013): Anterior insular cortex mediates bodily sensibility and social anxiety. Soc Cogn Affect Neurosci. 8: 259–266.

31. Dennis EL, Jahanshad N, McMahon KL, de Zubicaray GI, Martin NG, Hickie IB, et al. (2014): Development of Insula Connectivity Between Ages 12 and 30 Revealed by High Angular Resolution Diffusion Imaging. Hum Brain Mapp. 35: 1790–1800.

32. Lord C, Rutter M, DiLavore P, Risi S, Gotham K, Bishop S (2012): Autism Diagnostic Observation Schedule, second edition (ADOS-2) manual (Part I): Modules 1-4. Torrance, CA: Western Psychological Services.

33. Constantino JN, Gruber CP (2012): Social responsiveness scale (SRS). Western Psychological Services Torrance, CA.

34. Harrigan RL, Yvernault BC, Boyd BD, Damon SM, Gibney KD, Conrad BN, et al. (2016): Vanderbilt University Institute of Imaging Science Center for Computational Imaging XNAT: A multimodal data archive and processing environment. NeuroImage. 124: 1097–1101.

35. Asman AJ, Landman BA (2013): Non-local statistical label fusion for multi-atlas segmentation. Med Image Anal. 17: 194–208.

36. Ashburner J, Friston KJ (1999): Nonlinear spatial normalization using basis functions. Hum Brain Mapp. 7: 254–266.

37. Ashburner J, Neelin P, Collins DL, Evans A, Friston K (1997): Incorporating prior knowledge into image registration. NeuroImage. 6: 344–352.

38. Jenkinson M, Bannister P, Brady M, Smith S (2002): Improved optimization for the robust and accurate linear registration and motion correction of brain images. NeuroImage. 17: 825–841.

39. Smith SM (2002): Fast robust automated brain extraction. Hum Brain Mapp. 17: 143–155.

40. Woolrich MW, Ripley BD, Brady M, Smith SM (2001): Temporal autocorrelation in univariate linear modeling of FMRI data. NeuroImage. 14: 1370–1386.

41. Power JD, Barnes KA, Snyder AZ, Schlaggar BL, Petersen SE (2012): Spurious but systematic correlations in functional connectivity MRI networks arise from subject motion. Neuroimage. 59: 2142–2154.

42. Beckmann CF, Jenkinson M, Smith SM (2003): General multilevel linear modeling for group analysis in FMRI. NeuroImage. 20: 1052–1063.

43. Woolrich M (2008): Robust group analysis using outlier inference. NeuroImage. 41: 286–301.

44. Woolrich MW, Behrens TEJ, Beckmann CF, Jenkinson M, Smith SM (2004): Multilevel linear modelling for FMRI group analysis using Bayesian inference. NeuroImage. 21: 1732–1747.

45. Farb NAS, Segal ZV, Anderson AK (2013): Attentional modulation of primary interoceptive and exteroceptive cortices. Cereb Cortex N Y N 1991. 23: 114–126.

46. Barrett LF, Simmons WK (2015): Interoceptive predictions in the brain. Nat Rev Neurosci. 16: 419–429.

47. Worsley KJ (2001): Statistical analysis of activation images. Ch 14, in Functional MRI: An Introduction to Methods. In: Jezzard P, Matthews PM, Smith SM, editors. New York, NY: Oxford University Press, pp 251–270.

48. Winkler AM, Ridgway GR, Webster MA, Smith SM, Nichols TE (2014): Permutation inference for the general linear model. NeuroImage. 92: 381–397.

49. Harris PA, Taylor R, Thielke R, Payne J, Gonzalez N, Conde JG (2009): Research electronic data capture (REDCap)--a metadata-driven methodology and workflow process for providing translational research informatics support. J Biomed Inform. 42: 377–381.

50. Holm S (1979): A Simple Sequentially Rejective Multiple Test Procedure. Scand J Stat. 6: 65–70.

51. Azur MJ, Stuart EA, Frangakis C, Leaf PJ (2011): Multiple Imputation by Chained Equations: What is it and how does it work? Int J Methods Psychiatr Res. 20: 40–49.

52. Buuren S van, Groothuis-Oudshoorn K (2011): mice: Multivariate Imputation by Chained Equations in R. J Stat Softw. 45: 1–67.

53. Telzer EH, McCormick EM, Peters S, Cosme D, Pfeifer JH, van Duijvenvoorde ACK (2018): Methodological considerations for developmental longitudinal fMRI research. Dev Cogn Neurosci, Methodological Challenges in Developmental Neuroimaging: Contemporary Approaches and Solutions. 33: 149–160.

54. Craig KD (2009): The social communication model of pain. Can Psychol Can. 50: 22–32.

55. Uddin LQ, Nomi JS, Hebert-Seropian B, Ghaziri J, Boucher O (2017): Structure and function of the human insula. J Clin Neurophysiol Off Publ Am Electroencephalogr Soc. 34: 300–306.

56. Lenard Z, Studinger P, Mersich B, Kocsis L, Kollai M (2004): Maturation of cardiovagal autonomic function from childhood to young adult age. Circulation. 110: 2307–2312.

57. Khalsa S, Rudrauf D, Tranel D (2009): Interoceptive awareness declines with age. Psychophysiology. 46: 1130–1136.

58. Murphy J, Geary H, Millgate E, Catmur C, Bird G (2017): Direct and indirect effects of age on interoceptive accuracy and awareness across the adult lifespan. Psychon Bull Rev. 1–10.

59. Fiene L, Ireland MJ, Brownlow C (2018): The Interoception Sensory Questionnaire (ISQ): A Scale to Measure Interoceptive Challenges in Adults. J Autism Dev Disord. 48: 3354–3366.

60. Garfinkel SN, Seth AK, Barrett AB, Suzuki K, Critchley HD (2015): Knowing your own heart: Distinguishing interoceptive accuracy from interoceptive awareness. Biol Psychol. 104: 65–74.

61. Murphy J, Millgate E, Geary H, Ichijo E, Coll M-P, Brewer R, et al. (2018): Knowledge of resting heart rate mediates the relationship between intelligence and the heartbeat counting task. Biol Psychol. 133: 1–3.

62. Mazurek MO, Vasa RA, Kalb LG, Kanne SM, Rosenberg D, Keefer A, et al. (2013): Anxiety, Sensory Over-Responsivity, and Gastrointestinal Problems in Children with Autism Spectrum Disorders. J Abnorm Child Psychol. 41: 165–176.

63. Failla MD, Moana-Filho EJ, Essick GK, Baranek GT, Rogers BP, Cascio CJ (2017): Initially intact neural responses to pain in autism are diminished during sustained pain. Autism Int J Res Pract. 1362361317696043.

64. Moore DJ (2015): Acute pain experience in individuals with autism spectrum disorders: a review. Autism Int J Res Pract. 19: 387–399.

65. Herbert BM, Muth ER, Pollatos O, Herbert C (2012): Interoception across Modalities: On the Relationship between Cardiac Awareness and the Sensitivity for Gastric Functions. PLOS ONE. 7: e36646.

66. Ring C, Brener J, Knapp K, Mailloux J (2015): Effects of heartbeat feedback on beliefs about heart rate and heartbeat counting: a cautionary tale about interoceptive awareness. Biol Psychol. 104: 193–198.

67. Desmedt O, Luminet O, Corneille O (2018): The heartbeat counting task largely involves non-interoceptive processes: Evidence from both the original and an adapted counting task. Biol Psychol. 138: 185–188.

68. Murphy J, Brewer R, Hobson H, Catmur C, Bird G (2018): Is alexithymia characterised by impaired interoception? Further evidence, the importance of control variables, and the problems with the Heartbeat Counting Task. Biol Psychol. 136: 189–197.

69. Zamariola G, Maurage P, Luminet O, Corneille O (2018): Interoceptive accuracy scores from the heartbeat counting task are problematic: Evidence from simple bivariate correlations. Biol Psychol. 137: 12–17.

70. Poquérusse J, Pastore L, Dellantonio S, Esposito G (2018): Alexithymia and Autism Spectrum Disorder: A Complex Relationship. Front Psychol. 9. doi: 10.3389/fpsyg.2018.01196.

71. Bird G, Cook R (2013): Mixed emotions: the contribution of alexithymia to the emotional symptoms of autism. Transl Psychiatry. 3: e285.

72. DeVille DC, Kerr KL, Avery JA, Burrows K, Bodurka J, Feinstein JS, et al. (2018): The Neural Bases of Interoceptive Encoding and Recall in Healthy Adults and Adults With Depression. Biol Psychiatry Cogn Neurosci Neuroimaging. 3: 546–554.

